# PACAP-PAC1 Receptor Inhibition is Effective in Models of Opioid-Induced Medication Overuse Headache

**DOI:** 10.1101/2021.12.05.471295

**Authors:** Zachariah Bertels, Elizaveta Mangutov, Kendra Siegersma, Alycia Tipton, Amynah A Pradhan

**Affiliations:** Department of Psychiatry, University of Illinois at Chicago

**Author notes:** Corresponding Author: Amynah A. Pradhan, 1601 W. Taylor Street (MC 912), Chicago IL, 60612.

**Keywords:** preclinical model, nitroglycerin, morphine, aura, cortical spreading depression

## Abstract

Opioids are regularly prescribed for migraine and can result in medication overuse headache and dependence. We recently showed that pituitary adenylate cyclase activating polypeptide (PACAP) is upregulated following opioid administration or in a model of chronic migraine. The goal of this study was to determine if PACAP was a link between opioid use and headache chronification. We tested the effect of PACAP-PAC1 receptor inhibition in novel models of opioid-exacerbated migraine pain and aura; and examined the co-expression between mu opioid receptor (MOR), PAC1, and PACAP in headache-associated brain and peripheral regions.

To model opioid exacerbated migraine pain, mice were injected daily with morphine (10 mg/kg) or vehicle for 11 days. On days 3,5,7,9, and 11 they also received the known human migraine trigger nitroglycerin (0.1 mg/kg) or vehicle. To model opioid exacerbated aura, mice were treated with vehicle or morphine twice daily for 4 days (20 mg/kg on days 1-3, 40 mg/kg on day 4), a well-established paradigm for causing opioid-induced hyperalgesia. On day 5 they underwent cortical spreading depression, a physiological correlate of migraine aura. The effect of the PAC1 inhibitor, M65 (0.1 mg/kg), was tested in these models. Fluorescent in situ hybridization was used to investigate the expression of MOR, PAC1, and PACAP.

Only mice treated with combined morphine and nitroglycerin developed chronic cephalic allodynia (n=18/group). M65 reversed this hypersensitivity (n=9/group). Morphine significantly increased the number of CSD events (n=8-9/group); and M65 decreased this exacerbation by morphine (n=8-12/group). PAC1 and/or PACAP were highly co-expressed with MOR, and varied by region (n=6/group). MOR and PACAP were co-expressed in the trigeminal ganglia, while MOR and PAC1 receptor showed near complete overlap in the trigeminal nucleus caudalis and periaqueductal gray. The cortex showed similar cellular co-expression between MOR-PACAP and MOR-PAC1.

These results show that opioids facilitate the transition to chronic headache through induction of PACAPergic mechanisms. Antibodies or pharmacological agents targeting PACAP or PAC1 receptor may be particularly beneficial for the treatment of opioid-induced medication overuse headache.

## Introduction

Opioid analgesics in current use (e.g. hydrocodone, oxycodone, meperidine) act primarily at the mu opioid receptor (MOR) and are still commonly prescribed for migraine [1, 2]. While opioids may provide acute relief, chronic use results in increased severity and progression of migraine from an episodic to a chronic state [3, 4]; a phenomenon known as medication overuse headache (MOH) [5]. Further, excessive use of prescription opioids by migraine patients is also a significant public health issue [6-8]. Over 50% of patients in a recent study were prescribed opioids at some point for headache [2], and a large scale epidemiological study published in 2020 found that 36% of migraine patients continued to use or keep on hand opioids for headache management [1]. In another study, opioids were administered in over half of all emergency room visits for migraine, and repeat visits to the emergency room was associated with opioid prescription [9]. Even after successful initial opioid withdrawal, 20-50% of patients relapsed within the first year, and most within the first 6 months [10]. The continued prescription of opioids for migraine puts a substantial number of patients at risk of developing MOH and potentially for prescription drug misuse and abuse [11, 12]. There is thus a desperate need to find effective therapies for opioid-induced MOH.

Pituitary adenylate cyclase-activating peptide (PACAP) has emerged as a therapeutic target for migraine [13-16]. This neuropeptide is found in two forms as a 38 amino acid peptide (PACAP38) and a truncated version, PACAP27 [17]. PACAP38 is the predominant form making up 90% of the circulating peptide and is found in both the peripheral and central nervous system [18]. PACAP38 can bind to one of three Gs G-protein coupled receptors (GPCRs), VPAC1, VPAC2, or PAC1 [19]. However, its affinity for PAC1 is 100-fold higher relative to the other two receptors {Harmar, 2012 #1685}. In clinical studies, direct infusion of PACAP produces headache in healthy subjects and migraine in migraine patients [21, 22]. Further strengthening the relationship between PACAP and migraine, PACAP38 is increased in the interictal phase of migraine patients [23]; and the migraine therapeutic, sumatriptan, was correspondingly associated with reduced PACAP38 levels [24].

Preclinical work also shows that PACAP produces migraine like phenotypes as infusion in rodents causes light aversive behavior [25, 26], meningeal dilation, and increased c-fos expression in the trigeminal nucleus caudalis (TNC) [25]. PACAP knockout mice also have reduced susceptibility to the effects of nitroglycerin (NTG), a known human migraine trigger commonly used to model migraine in rodents [25]. The PACAPergic system may play a distinct role in opioid-induced MOH. Our lab recently performed an unbiased large scale peptidomic study comparing mouse models of opioid induced hyperalgesia (OIH) and chronic migraine-associated pain [27]. PACAP was one of the few neuropeptides with altered expression in both models. In confirmation experiments, we found that the PAC1 inhibitor, M65, blocked the development of cephalic allodynia in a NTG model of migraine-associated pain, as well as in a model of OIH using chronic escalating doses of morphine [27]. PACAP may act as bridge between opioid use and migraine chronification.

Although PACAP has been investigated as a target for migraine, to the best of our knowledge it has not been considered for opioid-induced MOH; and one of the aims of the current study was to explore this role further. To date, most preclinical studies either model chronic migraine or OIH. To better reflect clinical MOH, we first developed 2 models of opioid-exacerbated migraine. We modified the commonly used NTG model of chronic migraine in which high doses of NTG result in the development of chronic allodynia [28-32]. We found that much lower doses of NTG did not produce chronic hypersensitivity unless combined with chronic morphine. Further we investigated the effect of morphine on a mechanistically distinct model of migraine - cortical spreading depression (CSD), a physiological correlate of migraine aura [33]. We found that repeated escalating doses of morphine also increased CSD events. We next tested the PAC1 inhibitor, M65 within these models of opioid exacerbated migraine pain and aura. M65 effectively reduced MOH-associated symptoms in both models. Finally, we investigated the cellular expression of the PACAPergic system relative to the mu opioid receptor (MOR) using in situ hybridization; and observed co-expression of MOR with PACAP or PAC1 in key headache processing regions. Together our results establish two new models of opioid induced MOH and demonstrate that inhibition of the PACAPergic system may be an feffective therapeutic strategy for this disorder.

## Materials and Methods

### Animals

Studies used adult male and female C57BL6/J mice (Jackson Laboratories, Bar Harbor, ME. USA). All mice were group housed with a 12hour-12hour light-dark cycle, in which lights were turned on at 07:00 and turned off at 19:00. Food and water were available ad libitum and mice weighed between 20-30 g. On test days weights were recorded and the experimenters were blinded. All experimental procedures were approved by the University of Illinois at Chicago Office of Animal Care and Institutional Biosafety Committee, in accordance with Association for Assessment and Accreditation of Laboratory Animal Care (AAALAC) International guidelines and the Animal Care policies of the University of Illinois at Chicago. All results are reported according to Animal Research: reporting of In Vivo Experiments (ARRIVE) guidelines. All animals were monitored continuously throughout experiments and no adverse effects were observed. All injections were in a volume of 10 ml/kg.

### Sensory Sensitivity Testing

At the beginning of each experiment a basal test for mechanical threshold was recorded, mice were counterbalanced into groups from this measurement. All sensory sensitivity tests were conducted in the same behavior room. The behavior room is separated from the vivarium and has low light (∼35-50 lux) and low noise conditions. All tests were performed during the light cycle between 08:00 and 17:00. The testing rack consisted of individual plexiglass boxes with a 4 oz paper cup in each box. Mice were habituated to the testing racks for 2 consecutive days before the initial test day. On test days mice were again habituated to the racks 20 minutes before the first test measurement. Mice were tested while in the paper cup. The periorbital region, caudal to the eyes and near the midline, was tested to assess cephalic mechanical thresholds. Manual von Frey hair filaments were used in an up-and-down method [34]. The filaments had a bending force ranging from 0.008g to 2g. The first filament used was 0.4g. Following the first filament if there was no response a heavier filament (up) was used, alternatively a response would result in the use of a lighter filament (down). Response were defined as shaking of the head, repeated pawing, or cowering away from the filament following a bend. Following the first response the up down method was continued for 4 additional filaments. Mice were tested with the PAC1 inhibitor, M65 (0.1 mg/kg IP); the CGRP receptor antagonist, olcegepant (1 mg/kg IP), or vehicle (saline) on day 12, 24h after their final morphine/NTG injection.

### Model of Opioid Exacerbated Migraine

Nitroglycerin (NTG) was purchased at a concentration of 5 mg/ml, in 30% alcohol, 30% propylene glycol and water (American Reagent, NY, USA). NTG was diluted on each test day in 0.9% saline to make a working solution of 1 mg/ml. This solution was further diluted in saline for a final dose of 0.01 mg/kg (0.001 mg/ml). Morphine (10 mg/kg) or vehicle (0.9% saline) was administered once daily subcutaneously for 11 days. Beginning on day 3, 30 minutes after the morphine/vehicle injection, animals received NTG (0.01 mg/kg, IP) or vehicle (saline). Basal mechanical thresholds were assessed on days 1, 3, 7, and 11 prior to that days’ morphine/vehicle treatment. A post-treatment measurement was also taken 2 hours after the NTG/Veh injection on days 3,7, and 11.

### Model of Opioid Exacerbated Aura

Before CSD measurement, mice underwent an OIH protocol as described previously [27, 35, 36]. Only female mice were used in this model. Briefly, mice received morphine (Sigma Chemical, St. Louis, MO) 20 mg/kg SC twice daily, once in the morning around 09:00 and again at 17:00, for days 1-3. On day 4 the dose was escalated to 40 mg/kg SC, which was also given twice that day. Vehicle, of 0.9% saline solution, was similarly administered to control mice. Mice were tested in the CSD model on day 5, 18-20h following their final morphine/vehicle injection. In a pilot experiment we also tested another morphine paradigm – 10 mg/kg SC daily for 11 days and CSD on day 12 – which did not significantly exacerbate CSD (Supplemental Data).

### Cortical Spreading Depression Model

The model of cortical spreading depression (CSD) used in these studies is based on previously published work [28, 37, 38]. Mice were randomly grouped into Vehicle or M65 (0.1 mg/kg, IP). For the CSD procedure mice were anesthetized with isoflurane (induction 3-4%; maintenance 0.75 to 1.25%; in 67% N2 / 33% O2) and placed in a stereotaxic frame on a homoeothermic heating pad. Core temperature (37.0 ± 0.5°C), oxygen saturation (∼ 99%), heart rate, and respiratory rate (80–120 bpm) were continuously monitored (PhysioSuite; Kent Scientific Instruments, Torrington, CT, USA). Mice were repeatedly tested with tail and hind paw pinch to ensure proper anesthetic levels were maintained.

CSD was verified in two ways, optical intrinsic signal (OIS) and electrophysiological recordings, as previously described [28, 38, 39] For OIS a thinned rectangular region of the skull was made about 2.5 × 3.3 mm^2^ (∼0.5 mm from sagittal, and ∼1.4 from coronal and lambdoid sutures) of the right parietal bone. A dental drill (Fine Science Tools, Inc., Foster City, CA, USA) was used to thin the skull. To increase transparency mineral oil was applied to the thinned region, which allowed for further visualization of the parenchyma and vasculature. For video recording a green LED (520 nm) light illuminated the skull throughout the experiment (1-UP; LED Supply, Randolph, VT, USA). Reflectance was collected with a lens (HR Plan Apo 0.5 × WD 136) through a 515LP emission filter on a Nikon SMZ 1500 stereomicroscope (Nikon Instruments, Melville, NY, USA). Images were acquired at 1–5 Hz using a high-sensitivity USB monochrome CCD (CCE-B013-U; Mightex, Pleasanton, CA, USA) with 4.65-micron square pixels and 1392 × 1040 pixel resolution.

Two burr holes were drilled lateral to the thinned window around the midpoint of the rectangle. The burr holes were drilled deeper than the thinned skull portion such that the dura was exposed, but not so deep that the dura was broken. Local field potentials (LFPs) were recorded using a pulled glass pipette filled with saline and attached to an electrode, which was further connected to an amplifier. The electrode was placed inside of the lateral burr hole such that it was inside of the cortical tissue. A separate ground wire was placed underneath the skin caudal to the skull, which was used to ground the LFPs. After set up, the LFP was recorded for an hour to allow for stabilization in the case that a CSD occurred during the placement of the electrode or the thinning surgery. After stabilization a second pulled glass pipette was filled with 1M KCl and placed into the rostral bur hole, ensuring there was no direct contact with the brain or surrounding skull. Once placed, an initial flow of KCl was started and an even flow was maintained so that a constant small pool of KCl filled the burr hole. Any excess liquid was removed with tissue paper. After initial KCl administration mice were recorded for 400 seconds. Following which mice were treated with M65 or vehicle and then recorded for a remaining 3600 seconds, for a total recording time of 4000 seconds. Animals were only included in the final analysis if at least 2 CSD events occurred within the first 400s of recording. No animals were excluded in this study. Following the recording, video and LFP were analyzed and used to count the number of CSD events that occurred within the hour recording. Following the procedure mice were euthanized by anesthetic overdose followed by decapitation.

### RNAScope Fluorescent In Situ Hybridization

RNAscope kit was purchased from Advanced Cell Diagnostics RNAScope Technology (ACD Bioscience). C57Bl6/J mice were anesthetized, brain and TG were collected and immediately frozen. Frozen tissue was cut on a cryostat at 14 µm, collected on slides, and processed per the manufacturer’s protocol. Every 4th section was quantified for each brain region. The probes used were targeted against the mouse genes for Oprm1, Adcyap1, and Adcyap1r1.

### Statistical Analysis

The sample size needed for each experiment was either based on similar previous experiments or calculated through power analysis where the minimal detectable difference in means =0.3, expected standard deviation of residuals= 0.2, desired power=0.8, alpha=0.05. Each experiment was replicated with separate cohorts of animals to ensure reproducibility. All data were analyzed using GraphPad Prism (GraphPad, San Diego, CA). The level of significance (α) for all tests was set to p<0.05. Unpaired, two-tailed, t-test was performed to determine effect of morphine on CSD; 2-way ANOVA to determine effect of M65 on morphine-CSD. Three-way ANOVA was performed for allodynia experiments. Post hoc analysis was conducted using Holm-Sidak analysis to correct for multiple comparisons. Post hoc analysis was only performed when F values achieved p < 0.05. All values in the text are reported as mean ± SEM.

## Results

### Repeated opioid administration exacerbates cephalic allodynia induced by low dose NTG

MOR agonists are still prescribed for the treatment of headache disorders despite the potential to cause MOH [4, 40, 41]. In this case opioids are overlaid on top of migraine pathophysiology, and our first aim was to establish migraine models that reflect this interaction. To this end we repeatedly treated animals with morphine (10 mg/kg, SC) or vehicle for 11 days and paired it with an intermittent low dose of the human migraine trigger NTG (0.01 mg/kg, IP) starting on day 3 (Figure 1A). We tested cephalic mechanical thresholds across the 11-day period; and responses were assessed before daily treatments (Basal Responses, Figure 1B), and 2h following the NTG/veh injection (Post-treatment Responses, Figure 1C). These doses of NTG or morphine alone did not produce chronic allodynia, but the combination of the two resulted in significant cephalic allodynia as observed on days 7 and 11 (Figure 1B). The low dose of NTG produced an acute allodynia 2 hours post-injection (Figure 1C, open squares). This hypersensitivity was blocked by morphine on the first day of NTG treatment (Figure 1C, filled square, day 3), but the anti-allodynic effects of morphine were lost by day 7. This data demonstrates that morphine tolerance is observed in this model, and that chronic opioid treatment exacerbates the effect of low dose NTG to produce chronic cephalic allodynia.

**Figure 1.**
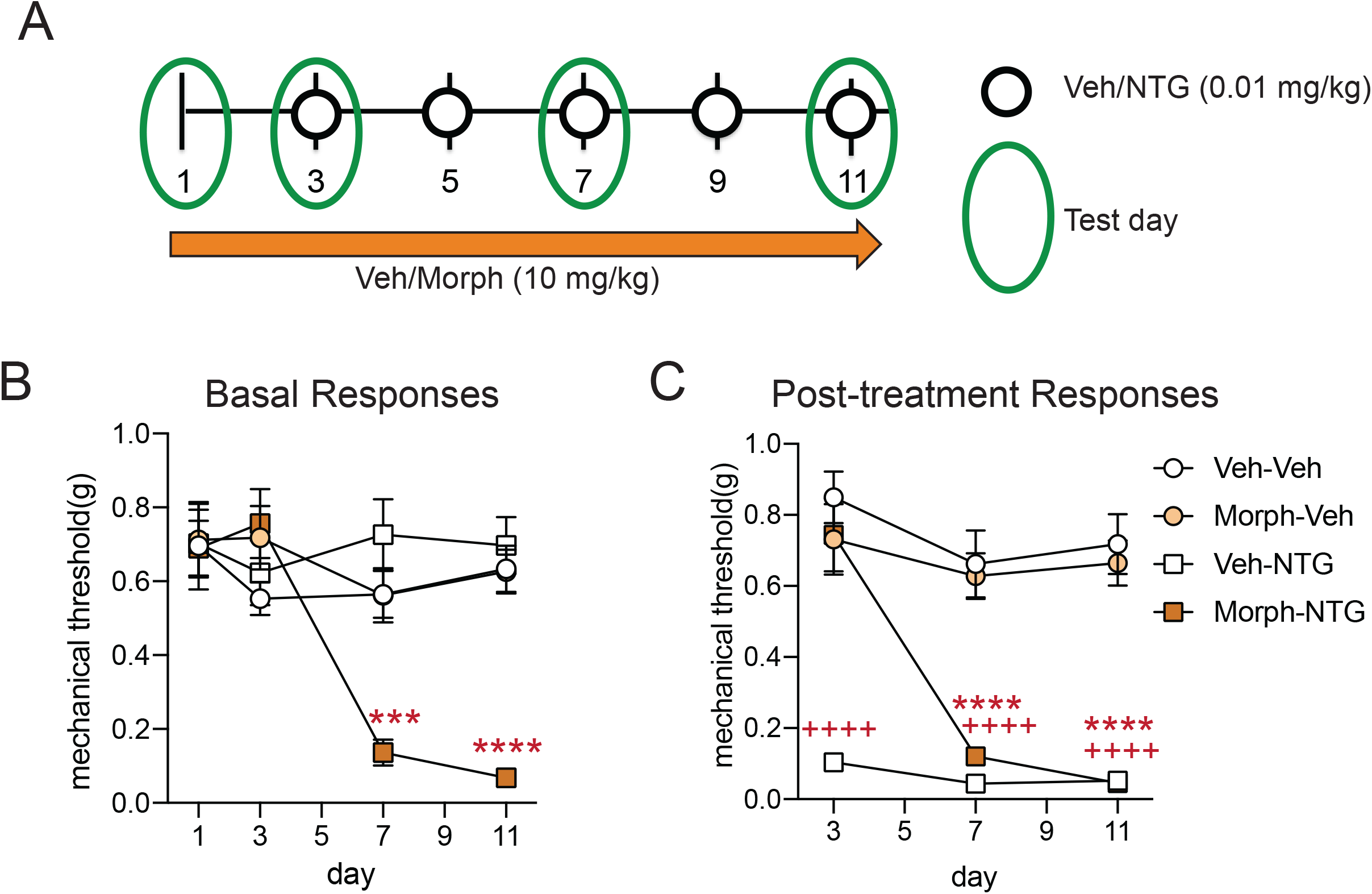
Morphine combined with low dose NTG induces chronic cephalic allodynia. (A) Schematic of testing and injection schedule, M&F C57BL6/J mice were treated every day with vehicle or morphine (10 mg/kg, SC; Morph) for 11 days. On days 3,5,7,9, and 11, mice were co-administered vehicle or low-dose nitroglycerin (0.01 mg/kg, IP; NTG) represented by white circles. The green oval shows days in which cephalic allodynia was assessed. (B) Periorbital mechanical thresholds were determined prior to drug treatment. Neither NTG nor morphine produced chronic allodynia alone. However, the combination of the two produced severe cephalic allodynia starting on day 7 which persisted to the end of treatment, p<0.001 effect of treatment, time, and interaction, three-way RM ANOVA and Holm-Sidak post hoc analysis. *** p<0.001; ****p<0.0001 relative to Veh-Veh on the same day, n=18/group. (C) Periorbital mechanical thresholds were also accessed 2 hours after vehicle, NTG, morphine administration to determine acute effects of treatment. On day 3, only the Veh-NTG group showed acute cephalic allodynia, but by day 7 the Morph-NTG groups also demonstrated severe allodynia. p<0.001 effect of treatment, time, and interaction, three-way RM ANOVA and Holm-Sidak post hoc analysis. ++++p<0.0001 Veh-NTG compared to Veh-Veh on same day. ****p<0.0001 Morph-NTG compared to Veh-Veh on same day n=18/group.

### Opioid induced hyperalgesia model results in increased cortical spreading depression

Approximately one third of migraine patients experience migraine aura as part of their symptoms, and cortical spreading depression (CSD) is thought to be the electrophysiological correlate of aura [33]. To examine if chronic MOR agonist treatment results in increased susceptibility to CSD we subjected mice to an established OIH model shown to cause cephalic allodynia [37, 42-45]. Mice were treated with morphine or vehicle twice daily for 4 days (Figure 2A, 20 mg/kg/injection on days 1-3 and 40 mg/kg/injection on day 4). As reported previously [37, 44, 45], this dosing regimen produced significant cephalic allodynia (Figure 2B). We tested mice in the CSD model [28, 37, 39, 46] on day 5, ∼16 hours after the final morphine treatment. Mice were recorded for an hour during continual administration of 1M KCl directly onto the dura. Total number of CSD events were counted based on visual reflectance changes (Figure 2C) and local field potential (LFP) recordings (Figure 2D). Chronic opioid treatment significantly increased CSD events relative to vehicle treated controls (Figure 2E). We also tested the dosing regimen of morphine used in the combined opioid-NTG model described in Figure 1. In this case, mice were treated with vehicle or morphine (10 mg/kg SC) daily for 11 days and CSD was tested on day 12. This dosing regimen of morphine does not cause cephalic allodynia (Figure 1B,C), and did not significantly exacerbate CSD (Supplemental Figure 1). These data indicate that OIH not only increases cephalic hyperalgesia but also increases frequency of CSD events.

**Figure 2.**
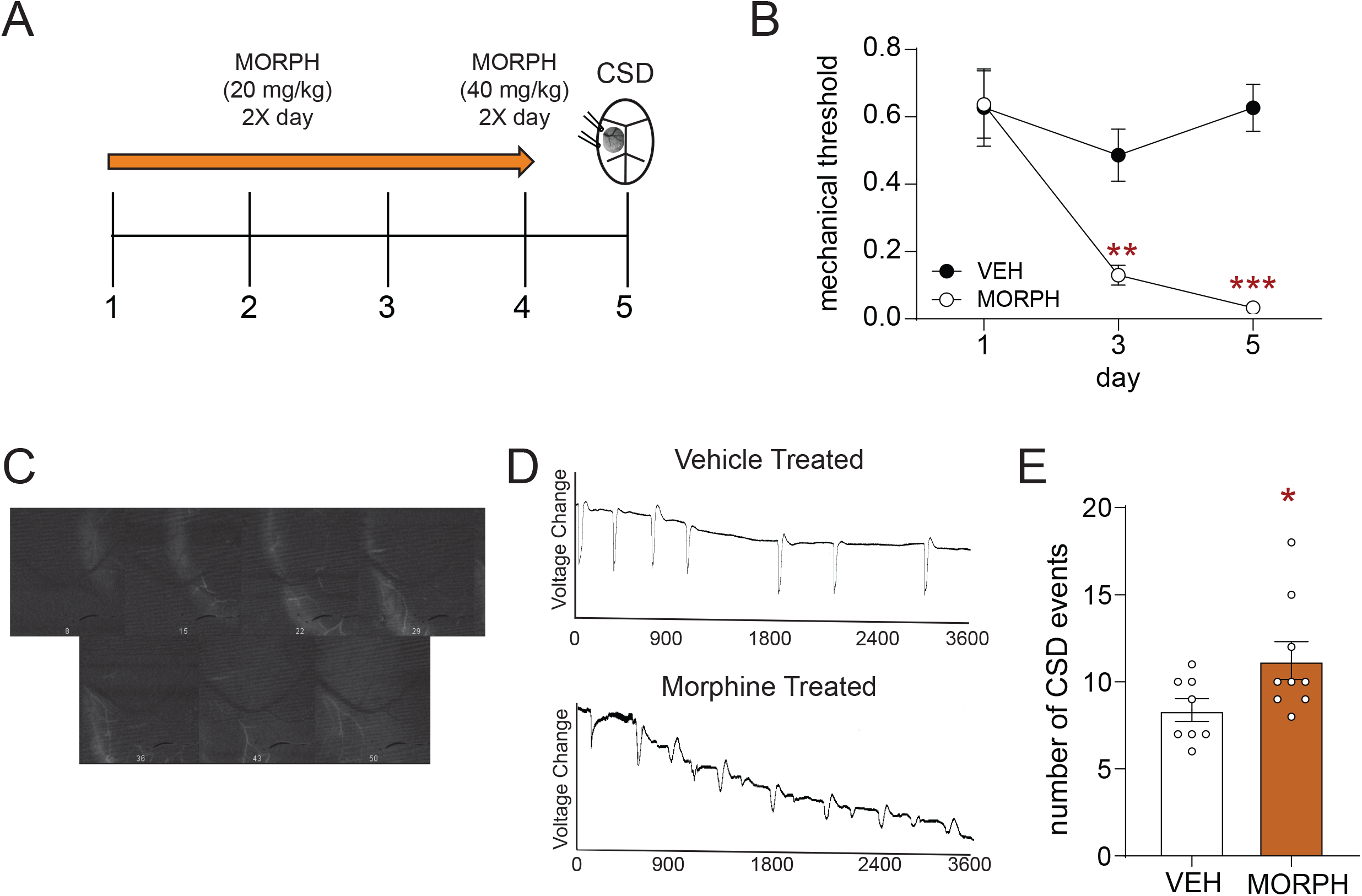
Chronic opioid treatment results in increased cortical spreading depression events. (A) Schematic of testing and injection schedule. Female C57BL6/J mice were treated twice daily with vehicle or morphine (20 mg/kg/injection Days 1-3, 40 mg/kg/injection Day 4, SC). On day 5 mice underwent CSD testing. (B) Periorbital mechanical thresholds were accessed prior to morning treatment of vehicle or morphine. Chronic morphine resulted in increased periorbital allodynia p<0.001 effect of treatment, time, and interaction, two-way ANOVA and Holm-Sidak Post hoc analysis. ****p<0.0001 Morph compared to Veh on Day 1 n=12/group. (C) Image sequence demonstrating reflectance change associated with an individual CSD event. (D) Representative line tracing of CSDs in vehicle (Top) or morphine (Bottom) treated mice over the 1 hour recording session. (E) Mice treated with morphine showed a significant increase in the average number of CSD events recorded over an hour. Unpaired t-test. *p<0.05 n=8-9/group.

### PAC1 antagonist inhibits allodynia induced by the mixed morphine-NTG model

We next determined if inhibition of the PACAPergic system could block cephalic allodynia in our opioid-facilitated migraine model. Mice were treated in the opioid facilitated migraine model as described above (Figure 3A). On day 12, 24 hours, after their final drug treatment, mice were injected with vehicle or the PAC1 antagonist, M65 (Figure 3A, 0.1 mg/kg IP) [47] and tested for cephalic thresholds 30 minutes post-treatment. At this time point, mice treated with morphine and NTG continued to show significant allodynia (Figure 3B, morphine-NTG-vehicle). Acute treatment with M65 completely inhibited allodynia in this group (Figure 3B, morphine-NTG-M65). Importantly, we also determined the effect of olcegepant (1 mg/kg IP, 2h post-injection), a calcitonin gene related peptide (CGRP) receptor antagonist, in this model. Olcegepant did not significantly inhibit allodynia induced by chronic morphine plus NTG (Figure 3B, morphine-NTG-OLC). These data suggest that inhibition of the PACAPergic system specifically can ameliorate opioid-facilitated migraine.

**Figure 3.**
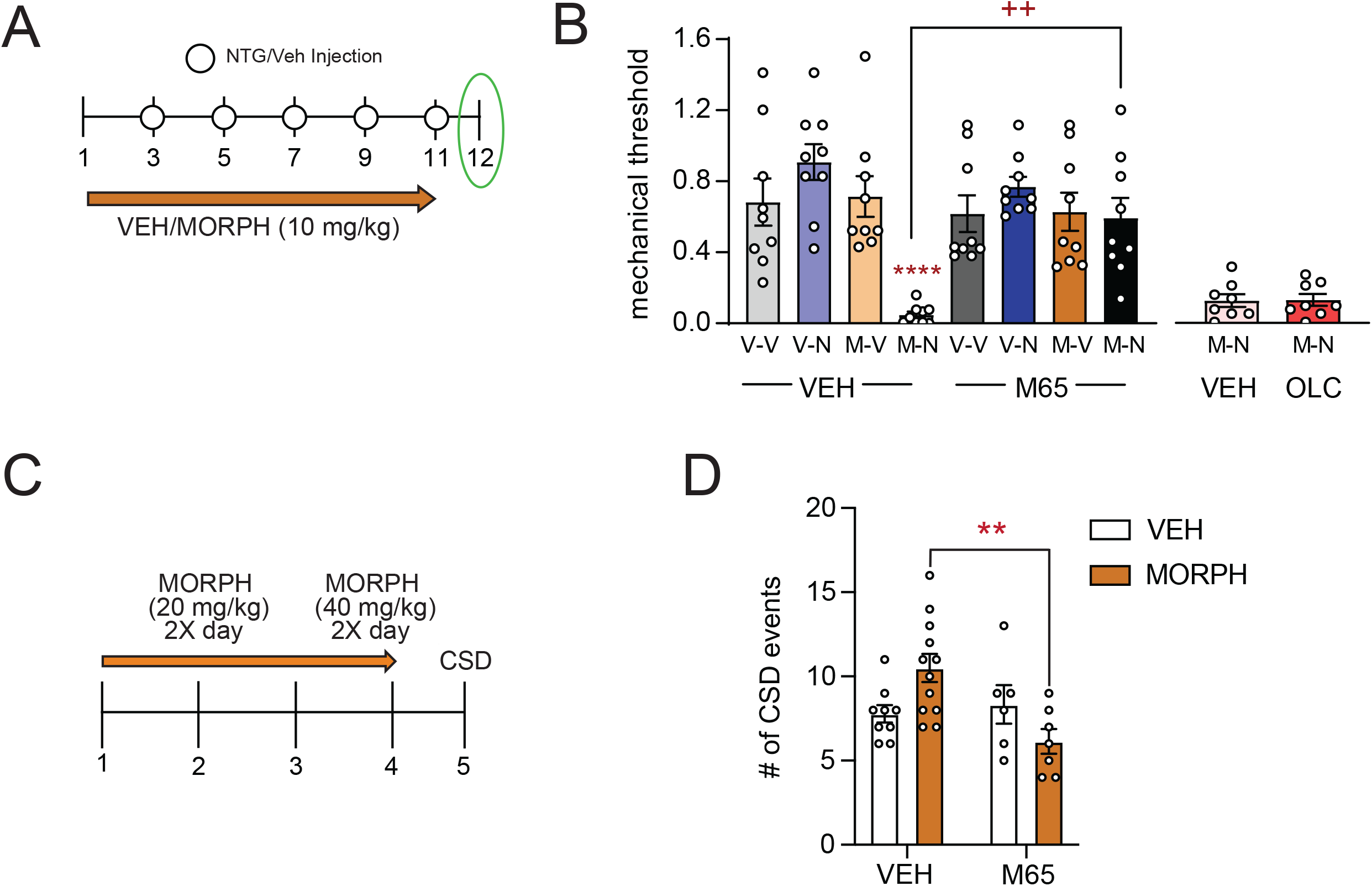
The PAC1 receptor inhibitor, M65, blocks opioid facilitation of NTG-induced allodynia and CSD. (A) Schematic of testing and injection schedule for model of opioid facilitated migraine pain. M&F C57BL6/J mice were treated every day with vehicle or morphine (10 mg/kg, SC; Morph) for 11 days. Starting on day 3, mice were co-administered vehicle or low-dose nitroglycerin (0.01 mg/kg, IP; NTG) for the remaining 9 days, represented by white circles. On day 12, 24 hrs follow the last treatment, mice were treated with either vehicle or M65 (0.1 mg/kg, IP) and tested 30 minutes later. (B) Periorbital mechanical thresholds were determined following M65 or vehicle administration. Chronic Morph-NTG treated mice continued to show cephalic allodynia at this time point; an effect that was inhibited by M65. p<0.001 interaction, three-way ANOVA and Holm-Sidak post hoc analysis. ****p<0.0001 Morph-NTG-Veh compared to Veh-Veh-Veh group; ++p<0.01 Morph-NTG-M65 compared to Morph-NTG-Veh. n=9/group. A separate group of mice were treated with morphine/NTG and then tested on day 12with vehicle or olcegepant (1 mg/kg IP, 2h post-injection). (C) Schematic of injection schedule for opioid facilitated CSD. Female C57BL6/J mice were treated twice daily with vehicle or morphine (20 mg/kg/injection days 1-3, 40 mg/kg/injection day 4, SC). On day 5 mice underwent CSD procedure. After the initial 400 seconds mice were treated with either vehicle or M65 (0.1 mg/kg IP) and recorded for an additional 3600 seconds. (D) Mice treated with chronic morphine showed an increased number of CSD events, and this increase was inhibited by M65 treatment. p<0.01 effect of M65, and interaction, two-way ANOVA and Holm-Sidak post hoc analysis. **p<0.01 Morph-Veh compared to Morph-M65 n=6-12/group

### PAC1 antagonist inhibits exacerbation of CSD by chronic morphine

We tested if PAC1 inhibition could affect CSD and/or opioid-exacerbation of CSD. Mice were treated in the 4 day OIH paradigm described above (Figure 3C), and on day 5 were tested in the CSD paradigm. Mice treated with vehicle or M65 (0.1 mg/kg IP) 400 seconds following the beginning of KCl stimulation and were recorded for an additional 3600 seconds. As was observed in Figure 2, chronic morphine resulted in a significant increase in CSD events in vehicle challenged mice (Figure 2D, left panel). Intriguingly, M65 did not reduce the number CSD events in chronic vehicle treat animals, but it significantly reduced the exacerbation of CSD by morphine (Figure 3D, right panel). These data suggest that a PACAPergic mechanism may mediate the facilitation of CSD by chronic opioid treatment.

### MOR is co-expressed with PACAP or PAC1 in migraine processing regions

We investigated if there was evidence for direct interaction between MOR and PACAP-PAC1 receptor. We used fluorescent in situ hybridization to map transcripts for *Oprm1* (MOR, yellow), *Adcyap1* (PACAP, magenta), and *Adcyap1R1* (PAC1, green); and examined the following migraine/pain processing regions: trigeminal ganglia (TG), trigeminal nucleus caudalis (TNC), somatosensory cortex (SSC), and lateral-ventrolateral periaqueductal gray (PAG) [48, 49] (Figure 4). In the TG, approximately 42% of MOR+ cells also expressed PACAP, while only ∼23% expressed PAC1. In contrast, in the TNC almost all MOR+ cells also co-expressed PAC1, while only ∼22% expressed PACAP. In the SSC, MOR was highly co-expressed with both PACAP and PAC1, ∼57% and ∼67%, respectively. Similarly, in the PAG we observed ∼50% co-expression of MOR and PACAP, and almost 100% co-expression of MOR+ cells with PAC1. These results demonstrate that there is very high cellular co-expression between MOR and components of the PACAPergic system.

**Figure 4.**
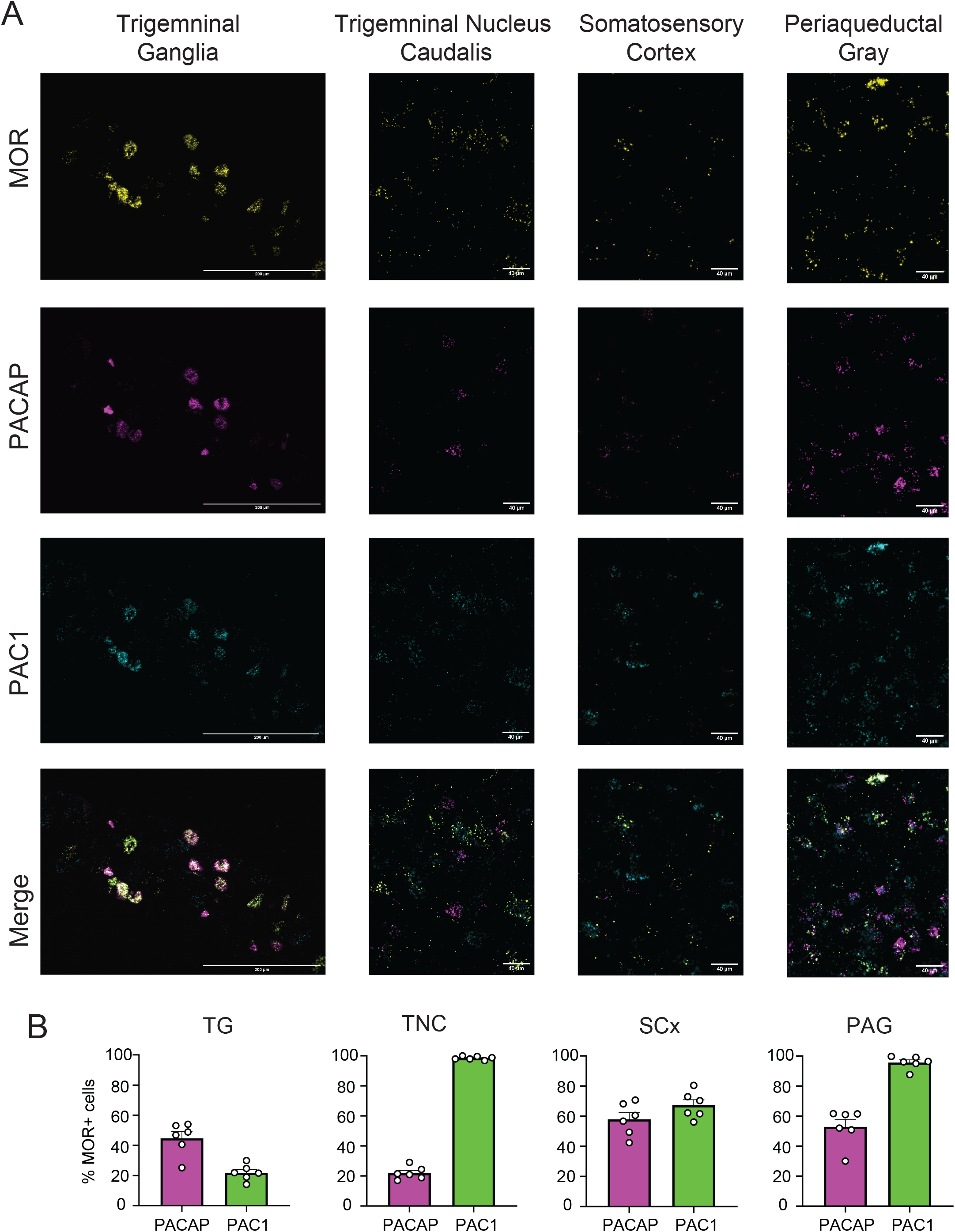
Co-expression of MOR with PACAP, and PAC1 in key migraine and pain processing regions. In situ hybridization was performed on brain slices from adult M&F mice using probes for *Oprm1* (MOR, yellow), *Adcyap1* (PACAP, magenta), and *Adcyap1R1* (PAC1, blue) (A) Representational images from trigeminal ganglia (TG), trigeminal nucleus caudalis (TNC), somatosensory cortex (SSC), and periaqueductal gray (PAG). (B) Percentage of MOR+ cells that co-express PACAP or PAC1 in these regions.

## Discussion

Opioids are still commonly prescribed for the treatment of headache disorders including migraine [1, 2, 40]. While opioids can provide limited relief from the acute migraine attack phase repeated use can result in transition to MOH [3, 4]. To better understand the drivers of opioid-induced MOH we developed two novel models which reflect opioid exacerbation of migraine-associated pain and aura. Chronic morphine treatment combined with a low dose of the human migraine trigger, NTG, resulted in chronic cephalic allodynia not observed with either treatment alone. We also demonstrated that a traditional OIH paradigm not only resulted in cephalic allodynia but also increased CSD events. We identify PAC1 receptor as a novel therapeutic target for opioid-induced MOH, as PAC1 inhibition blocked opioid facilitation in both of these models. Importantly, CGRP blockade was ineffective in reducing opioid-induced MOH. This finding suggests that opioid-induced MOH is distinctly regulated by the PACAPergic system; and not just by migraine mechanisms more generally. This study also demonstrates that MOR, PACAP, and PAC1 are co-expressed in numerous sites associated with headache processing, further supporting the idea of direct interaction between these two systems.

Clinically, OIH is treated by opioid taper and cessation, which can be difficult to implement as patients are reluctant to stop using drugs which they believe are treating their pain, and because of opioid withdrawal [50]. Even after successful initial opioid withdrawal, 20-50% of headache patients relapse within the first year, and most of those patients relapse within the first 6 months [51]. A greater understanding of the pathophysiology underlying MOH would allow for discovery of more targeted treatments for this disorder. One of our aims was to produce novel preclinical models that reflected the mechanistic interactions between chronic opioid treatment and migraine pathology. NTG evokes migraine in migraine patients [29], and has been used as a human experimental model of migraine [52]. Our lab has used chronic intermittent administration of higher dose NTG (10 mg/kg) to model chronic migraine-associated pain [53, 54]. In the current study we found that a much lower dose of NTG (0.01 mg/kg) induced acute cephalic allodynia but did not result in chronic hypersensitivity. Similarly, frequent and escalating doses of morphine results in OIH [43, 55, 56], however, the regimen used in our mixed NTG-opioid model was insufficient to produce OIH. Only when low-dose NTG and morphine were combined did we observe the development of chronic cephalic allodynia. MOH patients show increased pain sensitivity in cephalic and extra-cephalic regions [57], and in epidemiological studies MOH is associated with increased pain severity and cutaneous allodynia [1, 3]. Therefore, the allodynia observed in this animal model of opioid-facilitated migraine pain reflects clinical observations and may be of translational significance.

CSD captures a mechanistically different aspect of migraine relative to the NTG model. CSD is an electrophysiological correlate of migraine aura and widely held to be the cause of aura symptoms [33]. Susceptibility to CSD has been linked to increased cortical excitability. Genetic knock-in models of familial/monogenic forms of migraine show increased susceptibility to CSD [58, 59]. Furthermore, multiple migraine preventives with diverse sites of action inhibit CSD [60, 61], and this model is used to screen novel preventives. We show that pretreatment of mice with an opioid paradigm shown to cause OIH [27, 37, 43, 45, 56], also increases the number of CSD events in response to KCl administration. Increased CSD events were also observed following chronic treatment with paracetamol [62], and chronic sumatriptan exposure resulted in a decrease in the stimulation threshold required to generate a CSD event [63]. Together with our findings, these studies show that the mechanisms underlying CSD are sensitive to medication overuse, and a combined drug-CSD model can be used to model MOH.

In a previous study we had identified PACAP as a potential bridge between chronic opioid treatment and migraine chronicity [27]. We followed up on these findings and demonstrated that PAC1 inhibition effectively blocked both the chronic allodynia induced by NTG-morphine treatment, as well as the facilitatory effects of morphine on CSD. The latter findings are particularly intriguing as M65 did not itself decrease CSD events (i.e. in vehicle treated mice), as has been demonstrated for migraine preventives [60, 61]. PAC1 inhibition only decreased the augmentation of CSD induced by chronic morphine. These results support the idea that PACAPergic mechanisms are specifically enhanced in response to chronic opioid treatment which can feed into migraine pathophysiology. A number of studies indicate that MOR, PACAP, and PAC1 are expressed at similar anatomical sites, including the PAG, trigeminal complex, and somatosensory cortex [13, 64]. Our in situ hybridization studies confirmed expression in these regions and show that at a cellular level MOR is abundantly co-expressed with PACAP and PAC1. A limitation of this study is that transcript expression does not always correspond with protein expression or functional interaction, and future studies will address these issues more specifically. Nevertheless, these novel findings are the first step to showing that chronic opioid action at MOR could directly impact PACAP and PAC1 expression and function.

Targeting PACAP and its receptor PAC1 is under investigation as a potential migraine therapy [13, 16]. A rodent specific PAC1 antibody was able to inhibit evoked nociceptive activity in rats [65], and we have shown that M65 can block the development of chronic migraine-associated pain induced by NTG [27]. AMG-301, a human monoclonal antibody targeting the PAC1 receptor, recently completed a Phase II clinical trial for migraine; but did not show efficacy over placebo [66]. Although disappointing, the antibody was well tolerated with minimal adverse effects. These results may point to the need to target PACAP, which can also bind to VPAC1 and VPAC2. An anti-PACAP antibody developed by Lundbeck may address this question. In addition, peripheral PAC1 inhibition may be insufficient for efficacy in migraine. In our peptidomic study PACAP levels were most altered in the PAG following chronic NTG or morphine treatment [27], supporting the role of central PACAPergic mechanisms in migraine and MOH. The work presented in the current study suggests that PAC1 may be a particularly effective target for opioid-induced MOH. M65 effectively blocked chronic cephalic allodynia induced by morphine-NTG treatment, an effect not observed with olcegepant. Both PACAP and CGRP are implicated in migraine mechanisms, and both evoke migraine in migraine patients [67]. Headache patients are heterogeneous and spontaneous migraines may be endogenously generated in each individual by different combinations of pro-migraine peptides and signaling molecules. Our findings suggest that opioid-induced MOH may be particularly weighted towards PACAPergic mechanisms; and that PACAP-PAC1 is a therapeutic target for this disorder.

## Abbreviations

CSD: cortical spreading depression
MOR: mu opioid receptor
NTG: nitroglycerin
OIH: opioid induced hyperalgesia
PACAP: pituitary adenylate cyclase activating peptide
PAC1: Pituitary adenylate cyclase-activating polypeptide type I receptor
PAG: periaqueductal gray
SSC: somatosensory cortex
TG: trigeminal ganglia
TNC: trigeminal nucleus caudalis
VPAC1 or 2: vasoactive intestinal peptide receptor type 1 or 2

## Figure Legends

**Supplementary Figure 1.**
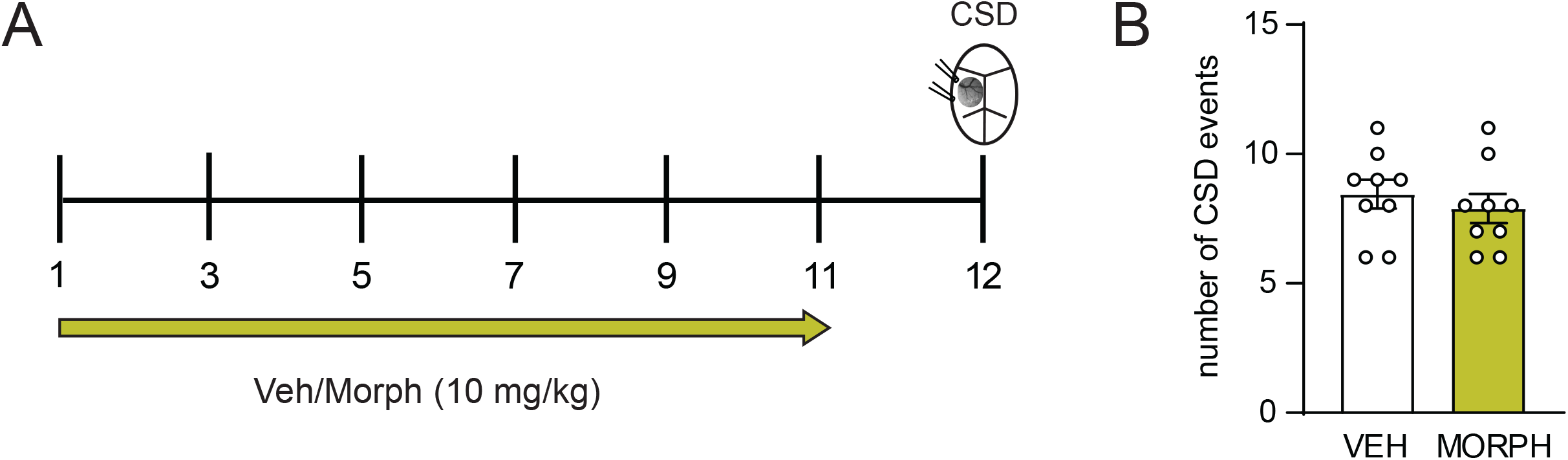
A protocol of morphine that does not induce allodynia did not exacerbate cortical spreading depression. (A)Schematic of injection schedule. Female C57BL6/J mice were treated every day with vehicle or morphine (10 mg/kg, SC; Morph) for 11 days. On day 12 mice underwent CSD testing. This regimen of morphine did not cause cephalic allodynia as shown in Figure 1. (B) There was no significant difference in the number of CSD events in vehicle vs morphine treated mice.

## References

1. Lipton, R.B., et al., Characterizing opioid use in a US population with migraine: Results from the CaMEO study. Neurology, 2020. 95(5): p. e457–e468.

2. Minen, M.T., et al., Survey of Opioid and Barbiturate Prescriptions in Patients Attending a Tertiary Care Headache Center. Headache, 2015. 55(9): p. 1183–91.

3. Schwedt, T.J., et al., Factors associated with acute medication overuse in people with migraine: results from the 2017 migraine in America symptoms and treatment (MAST) study. J Headache Pain, 2018. 19(1): p. 38.

4. Bigal, M.E. and R.B. Lipton, Excessive opioid use and the development of chronic migraine. Pain, 2009. 142(3): p. 179–182.

5. Headache Classification Committee of the International Headache Society (IHS) The International Classification of Headache Disorders, 3rd edition. Cephalalgia, 2018. 38(1): p. 1–211.

6. Colas, R., et al., Chronic daily headache with analgesic overuse: epidemiology and impact on quality of life. Neurology, 2004. 62(8): p. 1338–1342.

7. Reid, M.C., et al., Use of opioid medications for chronic noncancer pain syndromes in primary care. J.Gen.Intern.Med., 2002. 17(3): p. 173–179.

8. Diener, H.C., et al., Medication-overuse headache: risk factors, pathophysiology and management. Nat Rev Neurol, 2016. 12(10): p. 575–83.

9. Friedman, B.W., et al., Current management of migraine in US emergency departments: an analysis of the National Hospital Ambulatory Medical Care Survey. Cephalalgia, 2015. 35(4): p. 301–9.

10. Katsarava, Z., et al., Rates and predictors for relapse in medication overuse headache: a 1-year prospective study. Neurology, 2003. 60(10): p. 1682–1683.

11. Stovner, L., et al., The global burden of headache: a documentation of headache prevalence and disability worldwide. Cephalalgia, 2007. 27(3): p. 193–210.

12. Bigal, M.E., et al., Chronic migraine in the population: burden, diagnosis, and satisfaction with treatment. Neurology, 2008. 71(8): p. 559–566.

13. Vollesen, A.L.H., F.M. Amin, and M. Ashina, Targeted Pituitary Adenylate Cyclase-Activating Peptide Therapies for Migraine. Neurotherapeutics, 2018. 15(2): p. 371–376.

14. Akerman, S. and P.J. Goadsby, Neuronal PAC1 receptors mediate delayed activation and sensitization of trigeminocervical neurons: Relevance to migraine. Sci Transl Med, 2015. 7(308): p. 308ra157.

15. Waschek, J.A., S.M. Baca, and S. Akerman, PACAP and migraine headache: immunomodulation of neural circuits in autonomic ganglia and brain parenchyma. J Headache Pain, 2018. 19(1): p. 23.

16. Bertels, Z. and A.A.A. Pradhan, Emerging Treatment Targets for Migraine and Other Headaches. Headache, 2019. 59 Suppl 2: p. 50–65.

17. Miyata, A., et al., Isolation of a neuropeptide corresponding to the N-terminal 27 residues of the pituitary adenylate cyclase activating polypeptide with 38 residues (PACAP38). Biochem Biophys Res Commun, 1990. 170(2): p. 643–8.

18. Arimura, A., et al., Tissue distribution of PACAP as determined by RIA: highly abundant in the rat brain and testes. Endocrinology, 1991. 129(5): p. 2787–9.

19. Harmar, A.J., et al., International Union of Pharmacology. XVIII. Nomenclature of receptors for vasoactive intestinal peptide and pituitary adenylate cyclase-activating polypeptide. Pharmacol Rev, 1998. 50(2): p. 265–70.

20. Harmar, A.J., et al., Pharmacology and functions of receptors for vasoactive intestinal peptide and pituitary adenylate cyclase-activating polypeptide: IUPHAR review 1. Br J Pharmacol, 2012. 166(1): p. 4–17.

21. Birk, S., et al., The effect of intravenous PACAP38 on cerebral hemodynamics in healthy volunteers. Regul Pept, 2007. 140(3): p. 185–91.

22. Schytz, H.W., et al., PACAP38 induces migraine-like attacks in patients with migraine without aura. Brain, 2009. 132(Pt 1): p. 16–25.

23. Tuka, B., et al., Alterations in PACAP-38-like immunoreactivity in the plasma during ictal and interictal periods of migraine patients. Cephalalgia, 2013. 33(13): p. 1085–95.

24. Zagami, A.S., L. Edvinsson, and P.J. Goadsby, Pituitary adenylate cyclase activating polypeptide and migraine. Ann Clin Transl Neurol, 2014. 1(12): p. 1036–40.

25. Markovics, A., et al., Pituitary adenylate cyclase-activating polypeptide plays a key role in nitroglycerol-induced trigeminovascular activation in mice. Neurobiol.Dis., 2012. 45(1): p. 633–644.

26. Kuburas, A., et al., PACAP induces light aversion in mice by an inheritable mechanism independent of CGRP. J Neurosci, 2021.

27. Anapindi, K.D.B., et al., PACAP and other neuropeptides link chronic migraine and opioid-induced hyperalgesia in mouse models. Mol Cell Proteomics, 2019.

28. Pradhan, A.A., et al., delta-Opioid receptor agonists inhibit migraine-related hyperalgesia, aversive state and cortical spreading depression in mice. Br J Pharmacol, 2014. 171(9): p. 2375–84.

29. Iversen, H.K., J. Olesen, and P. Tfelt-Hansen, Intravenous nitroglycerin as an experimental model of vascular headache. Basic characteristics. Pain, 1989. 38(1): p. 17–24.

30. Christiansen, I., et al., Glyceryl trinitrate induces attacks of migraine without aura in sufferers of migraine with aura. Cephalalgia, 1999. 19(7): p. 660–667.

31. Olesen, J., The role of nitric oxide (NO) in migraine, tension-type headache and cluster headache. Pharmacol.Ther., 2008. 120(2): p. 157–171.

32. Olesen, J., H.K. Iversen, and L. Thomsen, Nitric oxide supersensitivity: a possible molecular mechanism of migraine pain. NeuroReport, 1993. 4(8): p. 1027–1030.

33. Charles, A.C. and S.M. Baca, Cortical spreading depression and migraine. Nat Rev Neurol, 2013. 9(11): p. 637–44.

34. Chaplan, S.R., et al., Quantitative assessment of tactile allodynia in the rat paw. J.Neurosci.Methods, 1994. 53(1): p. 55–63.

35. Liang, D.Y., et al., A genetic analysis of opioid-induced hyperalgesia in mice. Anesthesiology, 2006. 104(5): p. 1054–62.

36. Zhang, P., et al., Opioid-Induced Hyperalgesia Is Associated with Dysregulation of Circadian Rhythm and Adaptive Immune Pathways in the Mouse Trigeminal Ganglia and Nucleus Accumbens. Mol Neurobiol, 2019. 56(12): p. 7929–7949.

37. Dripps, I.J., et al., Forebrain delta opioid receptors regulate the response of delta agonist in models of migraine and opioid-induced hyperalgesia. Sci Rep, 2020. 10(1): p. 17629.

38. Bertels, Z., et al., A non-convulsant delta-opioid receptor agonist, KNT-127, reduces cortical spreading depression and nitroglycerin-induced allodynia. Headache, 2020.

39. Bertels, Z., et al., Neuronal complexity is attenuated in preclinical models of migraine and restored by HDAC6 inhibition. Elife, 2021. 10.

40. Buse, D.C., et al., Opioid use and dependence among persons with migraine: results of the AMPP study. Headache, 2012. 52(1): p. 18–36.

41. Mifflin, K.A. and B.J. Kerr, The transition from acute to chronic pain: understanding how different biological systems interact. Can J Anaesth, 2014. 61(2): p. 112–22.

42. Elhabazi, K., et al., Assessment of morphine-induced hyperalgesia and analgesic tolerance in mice using thermal and mechanical nociceptive modalities. J Vis Exp, 2014(89): p. e51264.

43. Chen, Y., C. Yang, and Z.J. Wang, Ca2+/calmodulin-dependent protein kinase II alpha is required for the initiation and maintenance of opioid-induced hyperalgesia. J Neurosci, 2010. 30(1): p. 38–46.

44. Anapindi, K.D.B., et al., PACAP and Other Neuropeptide Targets Link Chronic Migraine and Opioid-induced Hyperalgesia in Mouse Models. Mol Cell Proteomics, 2019. 18(12): p. 2447–2458.

45. Moye, L.S., et al., Delta opioid receptor agonists are effective for multiple types of headache disorders. Neuropharmacology, 2018. 148: p. 77–86.

46. Bertels, Z., et al., A non-convulsant delta-opioid receptor agonist, KNT-127, reduces cortical spreading depression and nitroglycerin-induced allodynia. Headache, 2021. 61(1): p. 170–178.

47. Uchida, D., et al., Maxadilan is a specific agonist and its deleted peptide (M65) is a specific antagonist for PACAP type 1 receptor. Ann N Y Acad Sci, 1998. 865: p. 253–8.

48. Goadsby, P.J., et al., Pathophysiology of Migraine: A Disorder of Sensory Processing. Physiol Rev, 2017. 97(2): p. 553–622.

49. Burstein, R., R. Noseda, and D. Borsook, Migraine: multiple processes, complex pathophysiology. J Neurosci, 2015. 35(17): p. 6619–29.

50. Hayhurst, C.J. and M.E. Durieux, Differential Opioid Tolerance and Opioid-induced Hyperalgesia: A Clinical Reality. Anesthesiology, 2016. 124(2): p. 483–8.

51. Katsarava, Z., et al., Rates and predictors for relapse in medication overuse headache: a 1-year prospective study. Neurology, 2003. 60(10): p. 1682–3.

52. Iversen, H.K., Human migraine models. Cephalalgia, 2001. 21(7): p. 781–5.

53. Pradhan, A.A., et al., Characterization of a novel model of chronic migraine. Pain, 2014. 155(2): p. 269–74.

54. Tipton, A.F., et al., The effects of acute and preventive migraine therapies in a mouse model of chronic migraine. Cephalalgia, 2016. 36(11): p. 1048–1056.

55. Ferrini, F., et al., Morphine hyperalgesia gated through microglia-mediated disruption of neuronal Cl(-) homeostasis. Nat Neurosci, 2013. 16(2): p. 183–92.

56. Hu, X., et al., PLGA-Curcumin Attenuates Opioid-Induced Hyperalgesia and Inhibits Spinal CaMKIIalpha. PLoS One, 2016. 11(1): p. e0146393.

57. Munksgaard, S.B., L. Bendtsen, and R.H. Jensen, Modulation of central sensitisation by detoxification in MOH: results of a 12-month detoxification study. Cephalalgia, 2013. 33(7): p. 444–53.

58. Eikermann-Haerter, K., et al., Abnormal synaptic Ca(2+) homeostasis and morphology in cortical neurons of familial hemiplegic migraine type 1 mutant mice. Ann Neurol, 2015. 78(2): p. 193–210.

59. van den Maagdenberg, A.M., et al., A Cacna1a Knockin Migraine Mouse Model with Increased Susceptibility to Cortical Spreading Depression. Neuron, 2004. 41: p. 701–710.

60. Bogdanov, V.B., et al., Migraine preventive drugs differentially affect cortical spreading depression in rat. Neurobiol.Dis., 2011. 41(2): p. 430–435.

61. Ayata, C., et al., Suppression of cortical spreading depression in migraine prophylaxis. Ann.Neurol., 2006. 59(4): p. 652–661.

62. Potewiratnanond, P., et al., Altered activity in the nucleus raphe magnus underlies cortical hyperexcitability and facilitates trigeminal nociception in a rat model of medication overuse headache. BMC Neurosci, 2019. 20(1): p. 54.

63. Green, A.L., et al., Increased susceptibility to cortical spreading depression in an animal model of medication-overuse headache. Cephalalgia, 2014. 34(8): p. 594–604.

64. Le Merrer, J., et al., Reward processing by the opioid system in the brain. Physiol Rev., 2009. 89(4): p. 1379–1412.

65. Hoffmann, J., et al., PAC1 receptor blockade reduces central nociceptive activity: new approach for primary headache? Pain, 2020. 161(7): p. 1670–1681.

66. Ashina, M., et al., A phase 2, randomized, double-blind, placebo-controlled trial of AMG 301, a pituitary adenylate cyclase-activating polypeptide PAC1 receptor monoclonal antibody for migraine prevention. Cephalalgia, 2021. 41(1): p. 33–44.

67. Ashina, M., et al., Human models of migraine -short-term pain for long-term gain. Nat Rev Neurol, 2017.

